# Cytoplasmic tail composition modulates the G protein and arrestin-3 signaling bias of the adhesion GPCR LPHN2

**DOI:** 10.1101/2025.07.04.663222

**Authors:** Krassimira Garbett, Chen Zheng, Julia Drube, Carsten Hoffmann, Vsevolod V. Gurevich, Richard C. Sando

## Abstract

The class B2 Adhesion GPCRs (aGPCRs) combine cell adhesion with GPCR signaling to control diverse developmental and physiological processes. How aGPCRs interact with and integrate distinct groups of effectors including G proteins, arrestins, and G protein receptor kinases (GRKs) remains unclear. Here, we find that diversity in the aGPCR C-terminal intracellular tail modulates G protein activation, arrestin-3 recruitment, and utilization of selective GRKs in LPHN2, a postsynaptic aGPCR essential for synapse formation. The C-terminal tail of LPHN2 is required for G protein activation and arrestin-3 recruitment. LPHN2 with an intact tail recruits arrestin-3 in the absence of G protein activation, suggesting constitutive arrestin-3-biased signaling. Alternative splicing of the LPHN2 tail modulates G protein activation and arrestin-3 binding independently, supporting that it controls G protein vs arrestin-3 bias. GRKs are important but not essential for arrestin-3 recruitment to LPHN2. Moreover, GRK2 increases arrestin-3 recruitment only in a subset of LPHN2 variants. Collectively, these results show that the mechanisms of the interactions of class B2 aGPCRs and arrestin are distinct from those of class A GPCRs and that splicing of the LPHN2 C-terminal tail determines G protein vs arrestin bias.

## Introduction

G protein-coupled receptors (GPCRs) are the largest family of signaling proteins in mammals. GPCRs integrate extracellular stimuli and direct appropriate cellular responses^1^. GPCR overstimulation is prevented by desensitization, which can last minutes (short-term) or hours (long-term)^2^. In general, short-term desensitization is achieved by phosphorylation of the GPCR by G protein-coupled receptor kinases (GRKs), followed by the binding of arrestins (arrestin1-4) that prevents further activation of cognate G proteins. The binding of arrestins also initiates internalization of the receptor via clathrin-mediated endocytosis, which leads to long-term receptor desensitization^3^. However, recent studies revealed that arrestins also control specific signaling pathways^4,5^, thereby serving as multifunctional adaptors and signal transducers^6–8^. GPCR signaling can be biased towards either G proteins or arrestin^9,10^. The cytoplasmic elements of GPCRs participate in G protein docking, are phosphorylated by GRKs and other kinases, and bind arrestins. Agonist-specific GPCR conformations may differentially affect the interactions with these three classes of binding partners thus generating signaling bias^11^. Dysregulation of GPCR signaling bias contributes to neuropsychiatric disorders, making it crucial to understand how different GPCR subfamilies achieve signaling selectivity.

Adhesion GPCRs (aGPCRs, GPCR family B2) are the second-largest GPCR subfamily and combine cellular adhesion with signal transduction. The extracellular region of aGPCRs is composed of several N-terminal domains involved in cell adhesion followed by a membrane proximal GAIN (GPCR Autoproteolysis-Inducing) domain. The GAIN is an evolutionarily conserved domain of aGPCRs that consists of ∼320 residues and has been a focus of aGPCR signaling mechanisms^12^. Many aGPCR GAIN domains contain an autoproteolytic cleavage site which generates a tethered agonist (TA), also known as the Stachel peptide^13,14^. Two models involving TA-induced signaling have been explored^15–17^. First, relatively strong adhesive force may remove the N-terminal fragment (NTF), exposing the TA and initiating a TA exposure-dependent downstream signaling. A second tunable mechanism involves the NTF remaining non-covalently bound to the C-terminal fragment (CTF) yet still signaling via the TA. Emerging evidence suggests that aGPCRs also use a third TA-independent mechanism, including conformational coupling between the extracellular NTF and 7-transmembrane (7TM) for transferring adhesion information to the GPCR core to modulate signaling^18–22^. Thus, context-specific extracellular interactions, once detected by the NTF of the aGPCR, are relayed inside the cell to initiate signaling. In contrast to class A GPCRs, only a few aGPCRs have been studied in relation to G protein and arrestin activation^23–28^. The relationship between different modes of TA-dependent or TA-independent activation mechanisms and arrestin recruitment, together with mechanisms of underlying bias towards G proteins and arrestins in aGPCRs, is largely unexplored.

Here we address the TA exposure-dependent mechanisms of G protein and arrestin-3 biased signaling by LPHN2 (ADGRL2), a postsynaptic aGPCR essential for synapse formation^29–31^. We demonstrate the contribution of specific cytoplasmic portions of the LPHN2 receptor towards G protein activation, arrestin-3 binding, and GRK preference by using mutant forms and splicing variants. Additionally, we reveal that the TA-exposure dependent arrestin-3 binding of LPHN2 is G protein-independent, and therefore likely represents an arrestin-3 signaling bias rather than receptor desensitization, and that alternative splicing of the C-terminus modulates this bias. Collectively, these results show that regulation of the aGPCR CTF composition in particular cell types likely fine-tunes signaling bias.

## Results

### aGPCRs that display TA exposure-dependent G protein activation recruit arrestin-3

aGPCR signaling can be both independent and dependent on TA exposure. Recent aGPCR engineering strategies allow for site-specific protease cleavage upstream of the TA and thus mimic TA exposure that may occur when strong adhesive force removes the aGPCR NTF^32,21^. We utilized an enterokinase (EK)-mediated proteolysis approach, which acutely exposes the native TA, to examine TA exposure-dependent arrestin-3 recruitment (**Fig. 1**). We replaced the NTF of six aGPCRs with critical neuronal functions – LPHN1-3 and BAI1-3 – with HaloTag-FLAG immediately upstream of the native TA sequence. These additional sequences allow the visualization of aGPCR fusions with HaloTag as well as EK-mediated cleavage following the FLAG sequence, which exposes the native TA peptide (**Fig. 1A**). We initially confirmed the surface localization of the six fusion proteins with a cell-impermeable HaloTag labeling protocol in HEK293T cells and found that all were trafficked to the plasma membrane (**Fig. 1B**).

**Figure 1:**
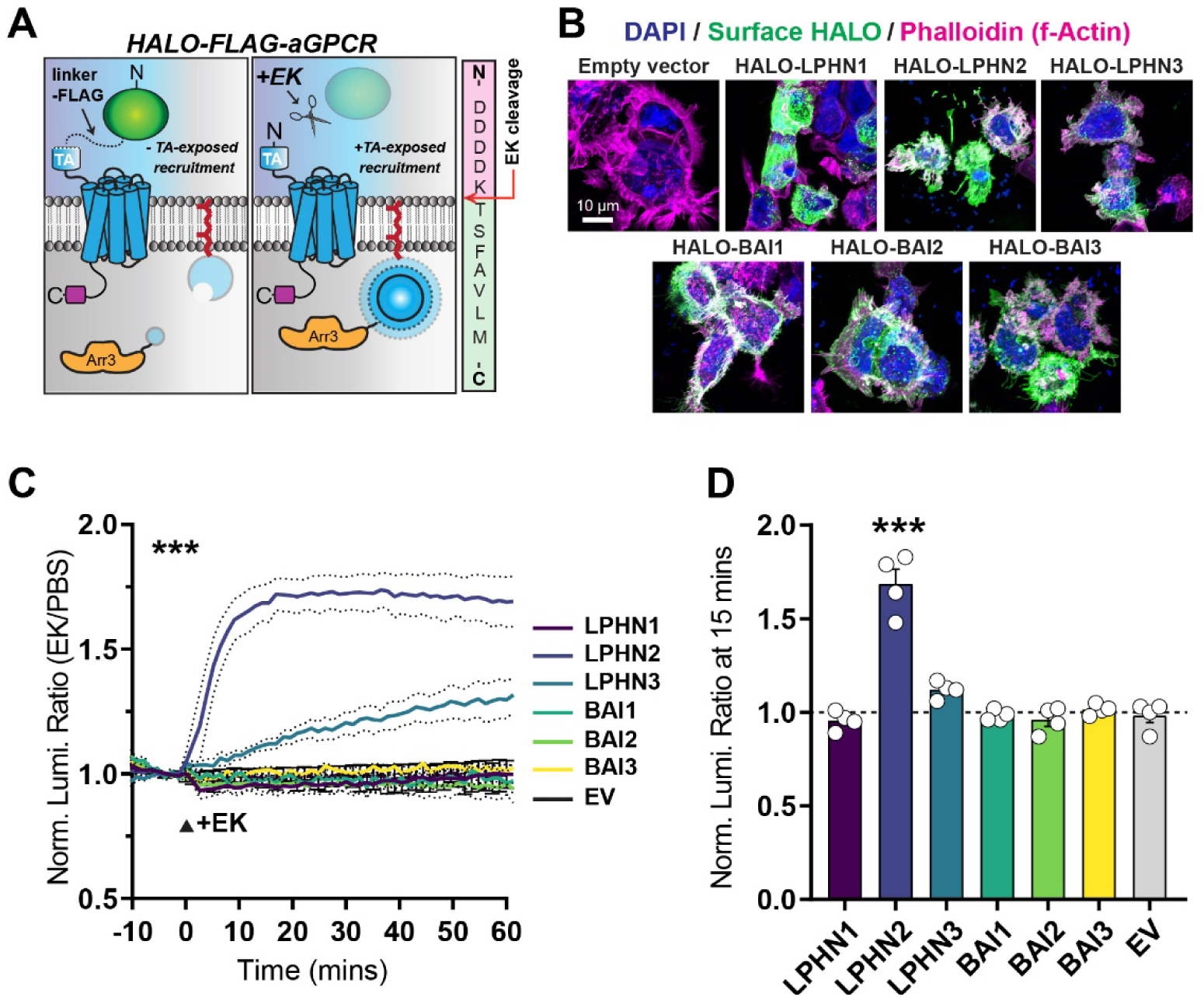
LPHN2 and LPHN3 recruit arrestin-3 upon TA exposure. **A,** diagram of experimental approach to test TA exposure-dependent arrestin-3 recruitment. Proteolysis via Enterokinase (EK) cleaves after the FLAG tag sequence, exposing the native tethered agonist (TA). **B,** representative images of surface HALO labeling of indicated receptor fusions. Labeling for DAPI and Phalloidin (f-Actin) was used as internal controls. **C,** NanoBiT arrestin-3 recruitment assay for LPHN1-3 or BAI1-3 compared to empty vector (EV). Luminescence was normalized to control (PBS addition). **D,** average normalized luminescence from NanoBiT assays in **C** at 15 min after EK treatment. Numerical data are means ± SEM from 4 independent biological replicates (depicted as open circles). Statistical significance was assessed with two-way ANOVA or one-way ANOVA with *post hoc* Tukey tests (***, p<0.001). See Figure S1 for additional arrestin-3 NanoBiT assays.

Next, we assessed TA exposure-dependent arrestin-3 recruitment using the NanoBiT complementation system, which uses Large BiT (LgBiT) and Small BiT (SmBiT) reconstitution to generate functional NanoLuc luciferase enzyme^33^. To avoid modification of the receptor, we expressed LgBit with the membrane-anchoring CAXX prenylation sequence. SmBit was fused at the arrestin-3 N-terminus, which does not participate in GPCR binding (**Fig. 1C & D**)^34^. The luminescence generated when LgBit and SmBit come together is indicative of arrestin-3 recruitment to the plasma membrane upon aGPCR activation. We conducted NanoBiT studies in arrestin-2/3 KO HEK293 cells to prevent potential interference of endogenous arrestins^35^. We measured baseline luminescence for 10 minutes, then treated the cells with EK to induce TA exposure and measured for an additional 60 minutes. Interestingly, of the six aGPCRs tested, only LPHN2, and LPHN3 to a lesser extent, displayed TA exposure-dependent arrestin-3 recruitment (**Fig. 1C & D**). While LPHN2 exhibited a typical logarithmic curve similar to the β2-adrenergic receptor (**Fig. S1**), LPHN3 TA exposure-dependent arrestin-3 membrane recruitment was slower, continuously increasing over the 60-minute period (**Fig. 1C & D**), suggesting that upon TA release, LPHN2 and LPHN3 engage arrestin-3 in different ways. Previous studies of these aGPCRs revealed that only LPHN2 and LPHN3 exhibit TA exposure-dependent G protein activation^36^, consistent with sequence predictions^37^. Thus, only a subset of aGPCRs demonstrate TA exposure-dependent signaling.

We next employed an arrestin-3 mutant, arrestin-3Δ7, to examine the molecular mechanisms of LPHN2/arrestin-3 recruitment (**Fig. 2**). The arrestin-3Δ7 has a 7-residue-deletion in its inter-domain hinge which prevents the binding to class A GPCRs^34,38^. We included β2-adrenergic receptor as a positive control, and empty vector as a negative control (**Fig. 2A & B**). As expected, the addition of 1 µM isoproterenol (ISO) induced robust arrestin-3 recruitment to β2-adrenergic receptor, which was substantially attenuated by the arrestin-3Δ7 mutation (**Fig. 2A & B**). The arrestin-3Δ7 mutation also reduced LPHN2 TA exposure-dependent arrestin-3 recruitment (**Fig. 2C & D**). However, at the end of the 60 minutes of measurement, a relatively higher response remained, suggesting that LPHN2 may interact with arrestin-3 in a different manner compared to well characterized class A GPCRs, such as β2-adrenergic receptor.

**Figure 2:**
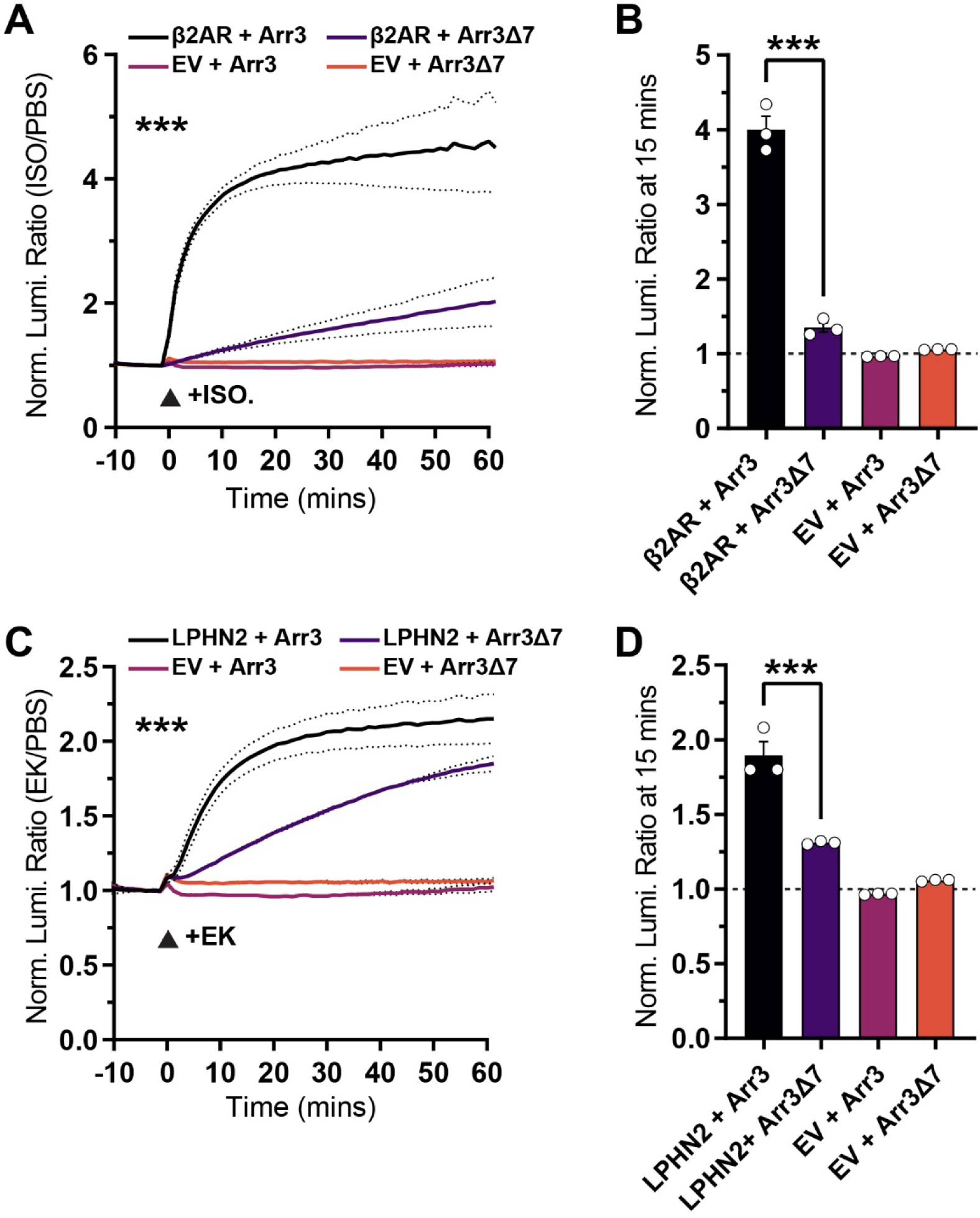
A hinge deletion arrestin-3 mutant reduces TA exposure-dependent arrestin-3 recruitment to LPHN2. **A,** validation of the arrestin-3Δ7 mutant with the β2-adrenergic receptor (β2AR) using arrestin-3 recruitment assay. **B,** average normalized luminescence from NanoBiT assay in **A** at 15 mins after 1 µM isoproterenol (ISO) treatment. **C,** arrestin-3 membrane recruitment assay with HALO-FLAG-LPHN2 and empty vector (EV) using SmBiT-arrestin-3 or the SmBiT-arrestin-3Δ7 mutant. **D,** average normalized luminescence from NanoBiT assay in **C** at 15 mins after EK treatment. Numerical data are means ± SEM from 3 independent biological replicates (depicted as open circles). Statistical significance was assessed with two-way ANOVA or one-way ANOVA with *post hoc* Tukey tests (***, p<0.001).

### ICL2 and the C-terminal cytoplasmic tail are required for LPHN2 TA exposure-dependent arrestin-3 recruitment

After receptor activation, arrestins bind to the phosphorylated C-terminus of many GPCRs, as well as to the receptor 7TM core^39,40^. The human LPHN2 variant used harbors a large (333 amino acids) cytoplasmic C-terminus with a PDZ-binding motif. Previous studies demonstrated that the LPHN C-terminus interacts with postsynaptic scaffolds including SHANKs and is essential for LPHN synaptic function^31,41,42^. Therefore, we tested the contribution of the LPHN2 7TM domain and C-terminus to TA exposure-dependent G protein activation and arrestin-3 binding by using a set of LPHN2 C-terminal and ICL2 mutants (**Fig. 3 & 4**). Like WT LPHN2, mutants had a HaloTag-FLAG at the N-terminus replacing the NTF to allow TA-exposure upon EK treatment. We generated a LPHN2 C-terminal tail truncation (LPHN2 ΔCt), a chimeric protein containing LPHN1 7TM and LPHN2 C-terminal tail (LPHN1-L2Ct), and a mutant containing two missense mutations V219A and F220A (LPHN2 AA) in intracellular loop 2 (ICL2) which prevent Gα12/13 coupling^43^. Mutant proteins were expressed, as detected by surface HaloTag labeling and immunoblotting (**Fig. 3B & C**), and trafficked to the membrane, like wild type LPHN2 (**Fig. 3B**).

**Figure 3:**
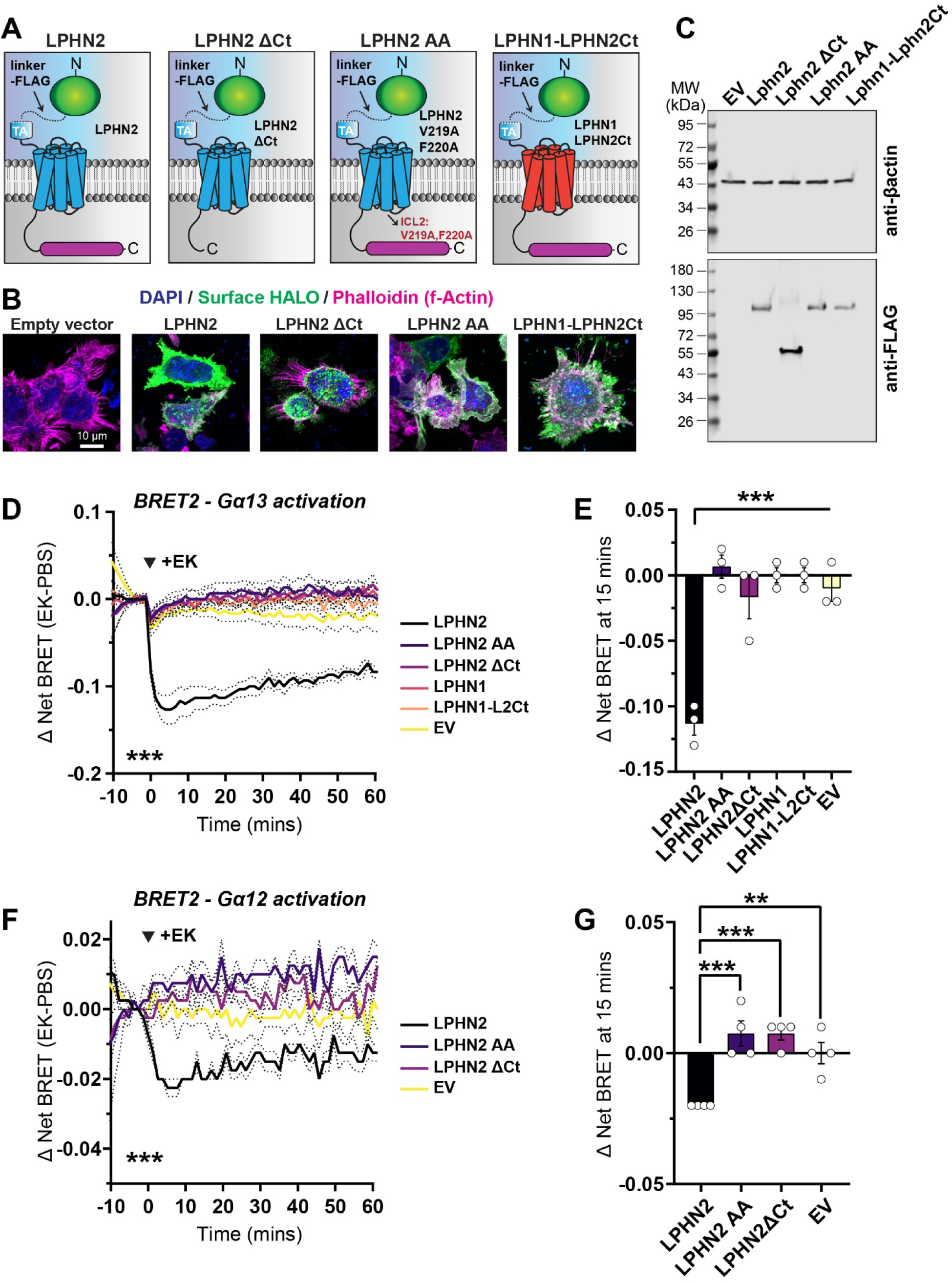
Intracellular loop 2 (ICL2) and the C-terminal cytoplasmic tail of LPHN2 are essential for TA exposure-dependent G protein activation. **A,** diagram of different human LPHN2 forms used for TRUPATH BRET2 G protein activation studies. **B,** representative images of surface HALO labeling of indicated receptors. Labeling for DAPI and (Phalloidin) f-Actin was used as internal controls. **C,** immunoblotting in indicated experimental conditions for FLAG tag. βactin served as a loading control. **D-G,** TRUPATH BRET2 assays for Gα13 (**D & E**) or Gα12 (**F & G**) in indicated conditions. **D,** time course of ΔNet BRET2 measurements using the Gα13 TRUPATH BRET2 sensor. Baseline BRET2 was measured for 10 mins and cells were subsequently treated with either EK or PBS at t = 0 and measured for an additional 60 mins. **E,** average ΔBRET2 (EK-PBS) measurements at 15 mins after treatment with EK or PBS. **F & G,** similar to **D** and **E**, except for experiments using the Gα12 TRUPATH BRET2 sensor. Numerical data are means ± SEM from 3-4 independent biological replicates (depicted as open circles). Statistical significance was assessed with two-way ANOVA or one-way ANOVA with *post hoc* Tukey tests (***, p<0.001; **, p<0.01).

We utilized TRUPATH BRET2 G protein activation sensors^44^ to determine the G protein activation efficacy of LPHN2 mutants in a G protein KO HEK293 cell line (GKO) to avoid interference of endogenous G proteins (**Fig. 3D-G**)^45^. Based on our previous demonstration that the human LPHN2 variant we used predominately activates Gα12/13 in a TA exposure-dependent manner^36^, we used the TRUPATH Gα12/13 BRET2 biosensors. We found that the deletion of the LPHN2 C-terminus abolished Gα12/13 activation (**Fig. 3D & E**). The LPHN1-L2Ct chimera also failed to activate G proteins like WT LPHN1. The LPHN2 AA mutation in ICL2 also impaired Gα12/13 activation, as reported previously^43^. Collectively, these results suggest that both ICL2 and the C-terminal tail are essential for G protein activation, and that neither of these LPHN2 elements alone is sufficient for coupling to Gα12/13.

### LPHN2 recruits arrestin-3 independently of G protein coupling

Next, we tested the ability of these LPHN2 mutants to recruit arrestin-3 (**Fig. 4**). LPHN2 ΔCt failed to recruit arrestin-3, suggesting that the C-terminal tail is essential for arrestin-3 binding (**Fig. 4A & B**). The LPHN1-LPHN2Ct chimera also failed to recruit arrestin-3. Surprisingly, while the LPHN2 AA mutant lacked TA-dependent G protein activation, it recruited arrestin-3 in a TA exposure-dependent fashion (**Fig. 4A & B**). The response was slower, but it reached the level of WT LPHN2 by 60 minutes. We next tested whether G proteins are required for arrestin-3 recruitment to LPHN2 by comparing WT and GKO HEK293 cells (**Fig. 4C & D**). LPHN2 TA exposure robustly induced arrestin-3 recruitment in GKO cells (**Fig. 4C & D**). Thus, LPHN2 with an intact C-terminus recruits arrestin-3 independently of G protein activation.

**Figure 4:**
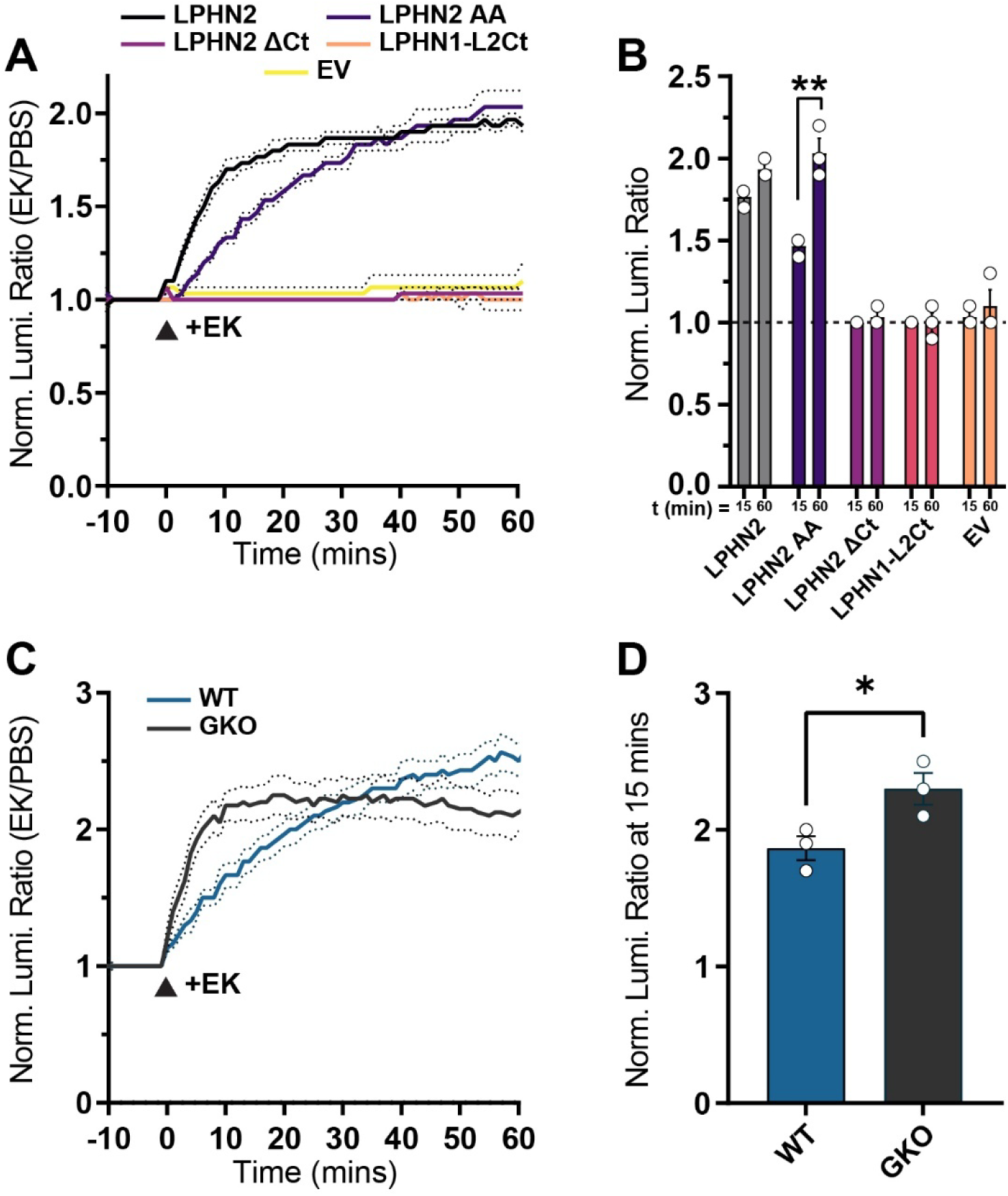
LPHN2 arrestin-3 recruitment requires the C-terminal tail but not G protein activation. **A,** arrestin-3 NanoBiT recruitment measurements in indicated experimental conditions. **B,** average normalized luminescence from NanoBiT assay in **A** at 15 mins and 60 mins after EK treatment. **C,** arrestin-3 NanoBiT membrane recruitment assay for HALO-FLAG-LPHN2 in either WT HEK293 or G protein KO (GKO) HEK293 cells. **D,** average normalized luminescence from NanoBiT assays in **C** at 15 mins after EK treatment. Numerical data are means ± SEM from 3 independent biological replicates (depicted as open circles). Statistical significance was assessed with two-tailed t-test (**, p<0.01; *, p<0.05).

### Alternative splicing of LPHN2 CTF differently affects G protein activation and arrestin-3 binding

Sequencing studies showed that LPHN1-3 are highly diverse in their exon usage and generate transcript variants that vary dramatically in their 7TM and C-terminal tail (**Fig. 5A & B**)^46^. We examined TA exposure-dependent G protein activation and arrestin-3 recruitment of eight CTF transcript variants compared to the previously used human LPHN2, which matches NCBI variant 1 (LPHN2v1) (**Fig. 5 & 6, Fig. S2**). We referred to the variants based on their NCBI transcript variant identifier for simplicity. The C-terminus length of selected variants (v1, v3, v4, v6, v7, v12, v13, v14, and v15) ranged from 34 to 366 amino acids due to alternate use of seven exons (**Fig. 5A**). LPHN2v13 had extra 16 amino acids due to incorporation of an additional exon that extended the ICL3 preceding TM6. We first ascertained cell surface expression of all variants (**Fig. 5C & D, Fig. S2**). We then examined G protein activation using the TRUPATH Gα12/13 BRET2 biosensors in GKO cells, normalizing responses to cell surface expression levels (**Fig. 5E & F, Fig. S2B**). None of tested variants had a higher TA-dependent Gα13 protein activation compared to LPHN2v1 (**Fig. 5E & F**). LPHN2v3, v6, v12, v14, and v15 demonstrated similar levels of Gα13 activation, while v4, v7, and v13 displayed diminished G protein activation compared to v1. Interestingly, v13 differs from v14 only by the 16 amino acid insertion in ICL3, suggesting that this insertion impairs TA exposure-dependent G protein coupling. Importantly, all variants displayed similar basal (TA exposure-independent) Gα12/13 activation, suggesting that splice variants differ only in TA exposure-dependent G protein signaling (**Fig. S3**).

**Figure 5:**
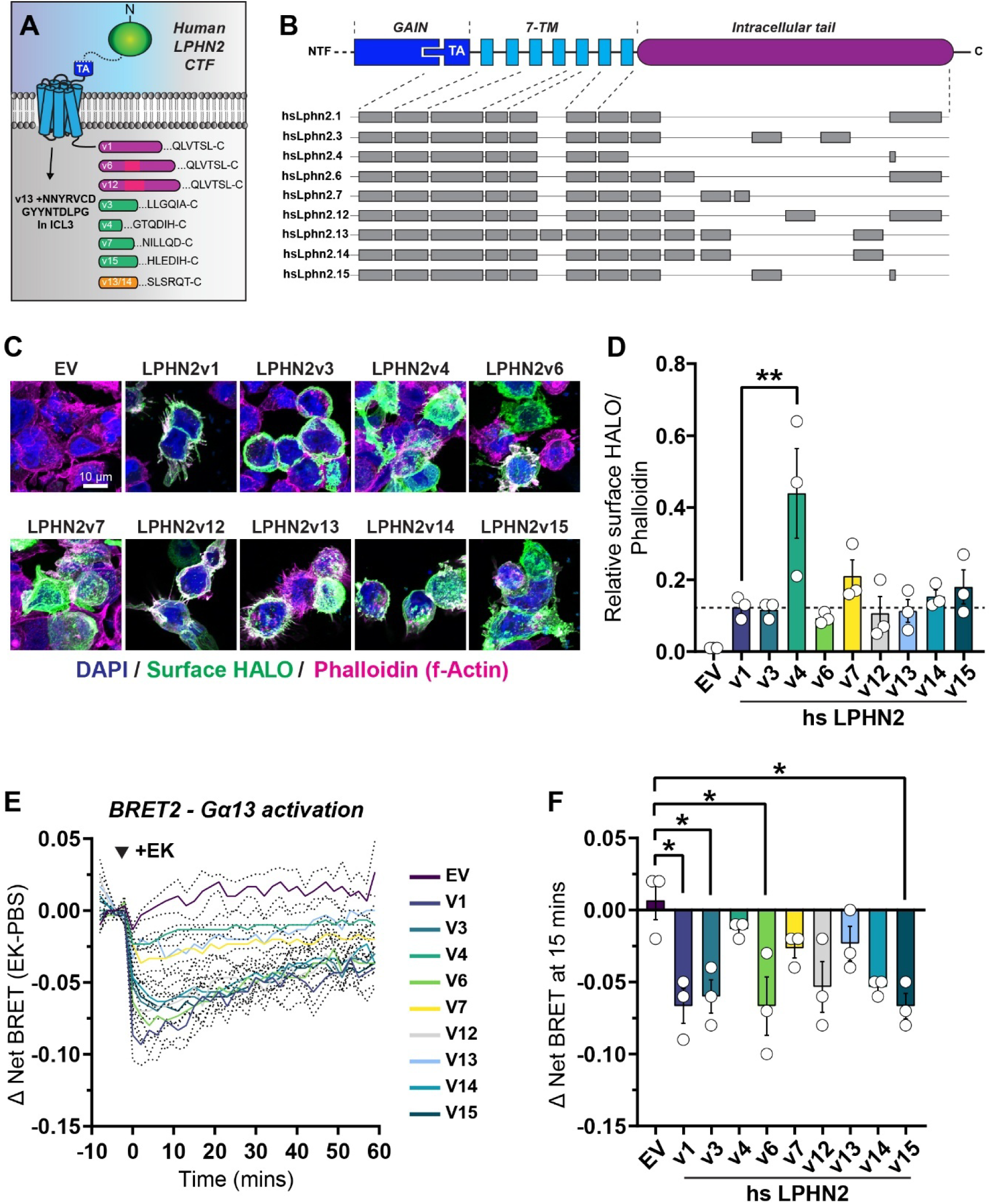
Human LPHN2 splice variants exhibit differential Gα12/13 activation efficacies. **A,** diagram of human LPHN2 transcript variants tested which display variation in intracellular loop 3 (ICL3) and the C-terminal cytoplasmic tail. **B,** diagram depicting the differential exon usage for each human LPHN2 variant. **C,** representative images of surface HALO labeling of indicated human LPHN2 transcript variants with N-terminal HALO-FLAG fusions. Labeling for DAPI and (Phalloidin) f-Actin was used as internal controls. **D,** quantification of HALO/Phalloidin signal ratios in indicated experimental conditions. **E,** time course of ΔNet BRET2 measurements using the Gα13 TRUPATH BRET2 sensor with indicated human LPHN2 variants and an empty vector (EV) control. **F,** average ΔBRET2 (EK-PBS) measurements at 15 mins after treatment with EK or PBS. Numerical data are means ± SEM from 3 independent biological replicates (depicted as open circles). Statistical significance was assessed with one-way ANOVA with *post hoc* Tukey tests (**. p<0.01; *, p<0.05). See Figure S2 and S3 for additional characterization of human LPHN2 variants.

We then assessed TA exposure-dependent arrestin-3 recruitment in arrestin-2/3 KO HEK293 cells (**Fig. 6**). Again, we normalized NanoBiT responses to the surface levels of expressed variants **(Fig. 5C & D).** Unexpectedly, several variants (v4, v7, v14, and v15) demonstrated higher TA-dependent arrestin-3 recruitment than v1, whereas LPHN2v3 and LPHN2v13 showed severely impaired arrestin-3 binding (**Fig. 6A & B**). Importantly, the magnitude of arrestin-3 recruitment did not correlate with G protein activation (**Fig. 5F, 6B & 6C**). For example, LPHN2v4 displayed relatively lower G protein activation yet enhanced arrestin-3 recruitment, while in terms of G protein activation levels LPHN2v3 was comparable to LPHN2v1, but its arrestin-3 recruitment was reduced (**Fig. 6A & B**). Additionally, the relatively small sequence difference between LPHN2v3 and LPHN2v7 was sufficient to reduce arrestin-3 recruitment to LPHN2v3 yet enhance it to LPHN2v7 relative to LPHN2v1. Thus, alternative splicing of the LPHN2 CTF directly affects the choice between G protein and arrestin-3 (**Fig. 6C**). Furthermore, the kinetics of arrestin-3 recruitment to splice variants were also different (**Fig. 6D, Fig. S4**). We fitted the corresponding curves to a logarithmic function and compared their slopes. Based on this parameter, LPHN2v1, v6 and v12 had similar kinetics, whereas LPHN2v4, v7, v14, and v15 demonstrated accelerated arrestin-3 binding (**Fig. 6D**). Thus, CTF sequence variations regulate the G protein/arrestin-3 bias and likely the desensitization kinetics.

**Figure 6:**
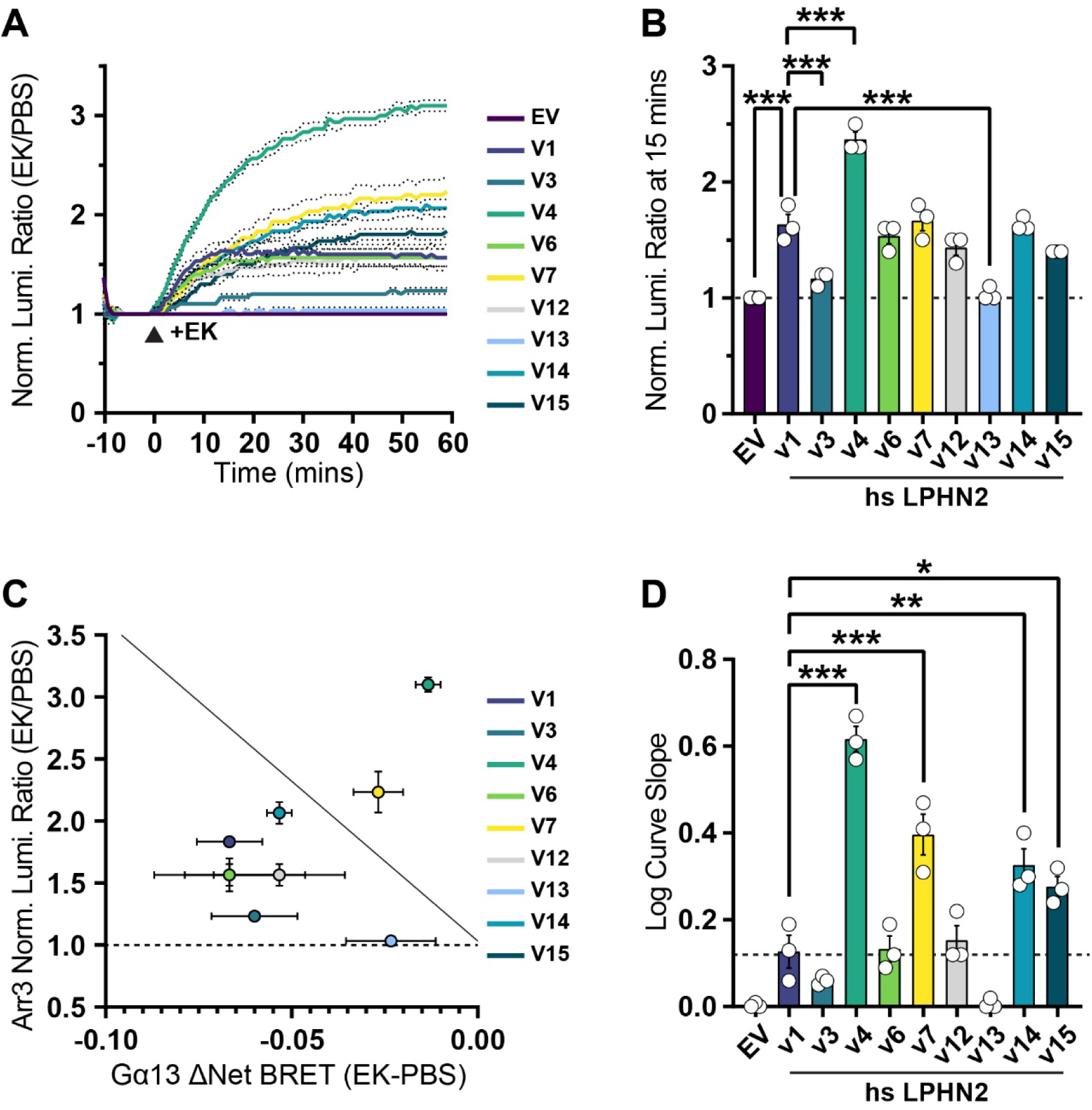
Variants in the LPHN2 CTF modulate arrestin-3 recruitment efficiency independently of G protein activation. **A,** time course of arrestin-3 NanoBiT recruitment in indicated experimental conditions. **B,** average normalized luminescence from NanoBiT assays in **A** at 15 mins after EK treatment. **C,** scatterplot depicting the correlation between average arrestin-3 recruitment and average Gα13 ΔNet BRET2 measurements of LPHN2 variants. **D,** quantification of the slope of the logarithmic curves of arrestin-3 recruitment. Numerical data are means ± SEM from 3 independent biological replicates (depicted as open circles). Statistical significance was assessed with one-way ANOVA with *post hoc* Tukey tests (***, p<0.001; **. p<0.01; *, p<0.05). See Figure S4 for additional information.

### GRKs contribute to LPHN2-arrestin-3 recruitment

The LPHN C-terminus contains numerous residues that can be phosphorylated. Therefore, we examined the role of GRKs in TA exposure-dependent arrestin-3 recruitment by LPHN2 (**Fig. 7**). To accomplish this, we generated a cell line lacking both arr2/3 and GRK2/3/5/6 using CRISPR/Cas9 (**Fig. S5**). We then compared arrestin-3 recruitment in arr2/3 KO and arr2/3 GRK2/3/5/6 KO HEK293 cells (**Fig. 7**). We used the β2-adrenergic receptor for comparison and found that agonist-dependent arrestin-3 recruitment was substantially attenuated in cells lacking GRK2/3/5/6, as expected (**Fig. 7A & B**)^47^. The loss of GRK2/3/5/6 also significantly impaired TA exposure-dependent arrestin-3 recruitment to LPHN2 (**Fig. 7C & D**). The residual recruitment in this cell line indicates that GRK2/3/5/6 are not absolutely required. We next determined the contribution of GRK2 towards arrestin-3 recruitment to different LPHN2 CTF variants by using arr2/3 GRK2/3/5/6 KO cells co-transfected with GRK2. Surprisingly, only a subset of LPHN2 variants relied on GRK2 (**Fig. 7E & F**). These included LPHN2 v4, v7, v14, v15, and v13 to a lesser extent. Interestingly, LPHN2 v1, v6, and v12, which appear to be GRK2-independent, have the largest C-terminal tail sharing the use of the final exon, which suggests that the shared sequence may inhibit the recruitment/activation of GRK2 upon TA release and/or contain serines/threonines phosphorylated by other kinases. Thus, GRK2-dependent arrestin-3 recruitment to LPHN2 upon TA release is CTF sequence-dependent.

**Figure 7:**
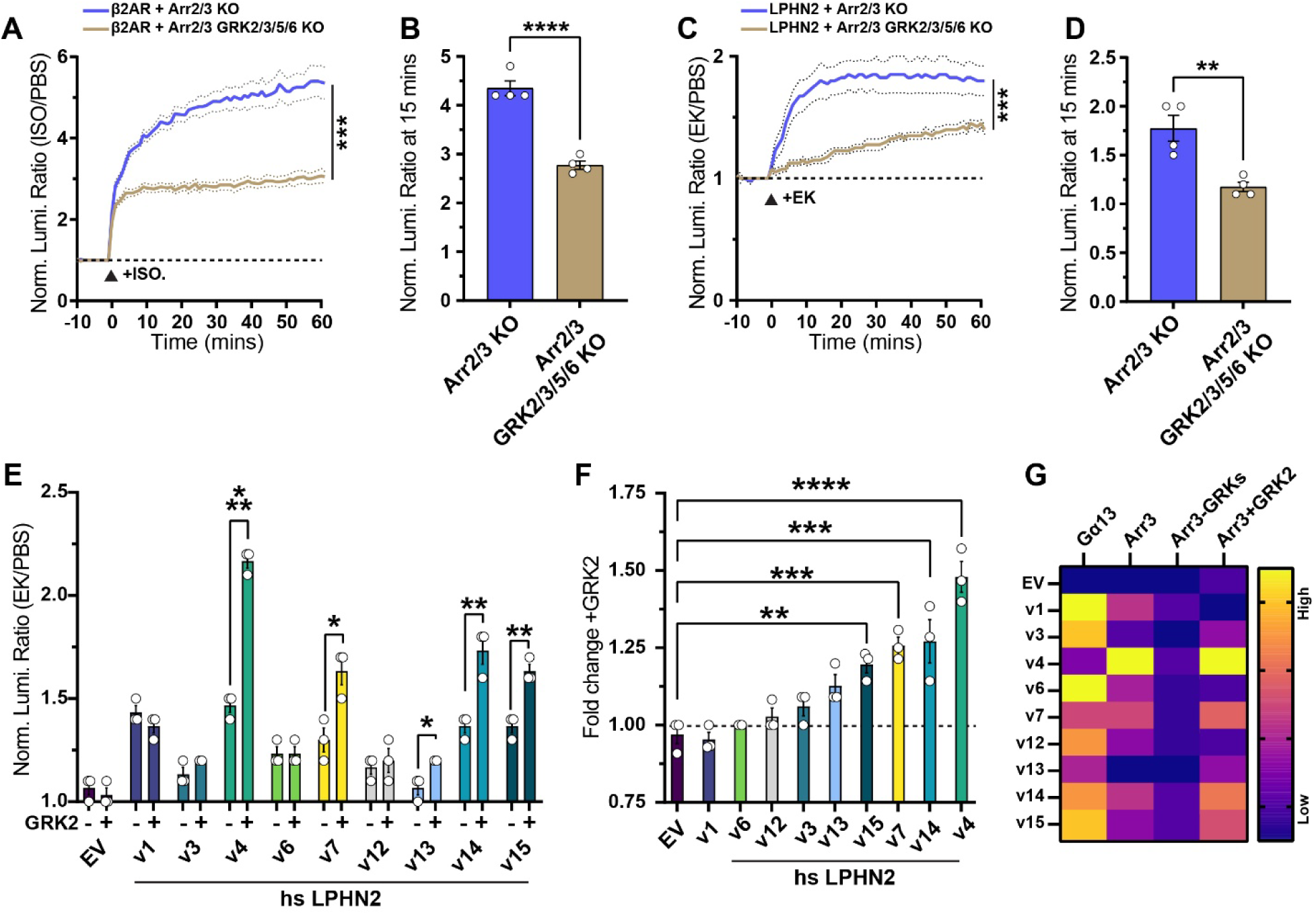
Loss of GRK2/3/5/6 attenuates arrestin-3 recruitment to LPHN2. **A & B,** arrestin-3 recruitment to β2AR in arr2/3 KO or arr2/3 GRK2/3/5/6 KO cells. **A,** time course of arrestin-3 recruitment. **B,** average normalized luminescence from NanoBiT assays in **A** at 15 mins after ISO treatment. **C & D,** similar to **A** and **B**, except for TA exposure-dependent LPHN2 arrestin-3 recruitment. **C,** time course of arrestin-3 recruitment. **D,** average normalized luminescence from NanoBiT assays in **C** at 15 mins after EK treatment. **E & F,** analysis of GRK2-dependent arrestin-3 recruitment by different LPHN2 CTF variants. **E,** arrestin-3 recruitment in arr2/3 GRK2/3/5/6 KO cells co-transfected with empty vector (-) or a GRK2 overexpression vector (+). **F,** fold change in luminescence shown in **E**. **G,** heat maps summarizing the results from G protein activation and arrestin-3 recruitment. The individual columns summarize the following: **Gα13** – average BRET2 TRUPATH Gα13 measurements; **Arr3** – average arrestin-3 NanoBiT recruitment measurements performed in arr2/3 KO HEK293 cells; **Arr3-GRKs** - average arrestin-3 NanoBiT recruitment measurements performed in arr2/3 GRK2/3/5/6 KO HEK293 cells; **Arr3+GRK2** – average fold-change difference of arrestin-3 NanoBiT recruitment measurements in arr2/3 GRK2/3/5/6 KO HEK293 cells overexpressing GRK2. Numerical data are means ± SEM from 3-4 independent biological replicates (depicted as open circles). Statistical significance was assessed with two-way ANOVA, one-way ANOVA with *post hoc* Tukey tests, or two-tailed t-test (***, p<0.0001; ***, p<0.001; **. p<0.01; *, p<0.05).

## Discussion

The postsynaptic aGPCRs LPHN1-3 integrate presynaptic ligand binding with intracellular signaling to control synapse assembly. Here, we investigated LPHN2 TA exposure-dependent G protein and arrestin-3 signaling bias. We found that a LPHN2 mutant that did not activate G proteins recruited arrestin-3. Furthermore, LPHN2 upon TA exposure robustly recruited arrestin-3 even in the absence of G proteins. Thus, LPHN2 does not require G proteins for arrestin-3 recruitment, which may indicate that LPHN2 has a major arrestin-3 signaling component. We showed that the variation of C-terminal tail sequences of LPHN2 by alternative splicing differentially affects G protein activation, GRK2 contribution, and arrestin-3 binding. Lastly, the accelerated kinetics of arrestin-3 recruitment to several LPHN2 splice variants appears to be GRK2-dependent, suggesting that, according to the barcode model^48,40,49^, these variants may initiate distinct signaling.

The preference of GPCRs for G proteins or arrestins depends on receptor sequence, bound ligands, and allosteric modulation^10^, leading to a multitude of intracellular responses^50^. Among the five GPCR subfamilies, the signaling bias of class B2 aGPCRs is perhaps the least understood. A distinguishing feature of the aGPCR subfamily is their large extracellular N-terminus containing multiple protein binding domains. Most aGPCRs also share the GAIN domain, which harbors an autoproteolysis site and TA. The activation of aGPCRs can occur by: (1) TA exposure via removal of the NTF, likely following strong mechanical force, (2) tunable signaling via the TA with an intact NTF, or (3) conformational coupling between the NTF and CTF in a TA-independent manner. Signaling bias of aGPCRs may be allosterically modulated, as observed for other GPCRs^51^, although this has not been explored. Here we utilized a proteolytic cleavage approach that leads to TA exposure. While we examined G protein and arrestin-3 interaction following TA exposure, it is tempting to speculate that aGPCRs may exhibit differential G protein vs arrestin-3 bias based on the mode of activation. In turn, context-specific combinations of adhesion partners and ligands present may promote a specific mode of activation. This versatility would increase the diversity of signaling events elicited by a given aGPCR. Experiments involving manipulation of each of the three modes of aGPCR activation are required to address this hypothesis.

Several recent structural studies with aGPCRs revealed the interactions of the TA with 7TM domain and their role in initiating G protein signaling^52–55^, but little is known about the similarities/differences between class A GPCRs and the aGPCRs in relation to arrestin recruitment^23^. Our results with the arrestin-3Δ7 mutant suggest that LPHN2 may interact with arrestins in a distinct manner compared to the class A β2-adrenergic receptor. Therefore, we explored what determines TA exposure-dependent G protein activation and arrestin-3 recruitment. The cytoplasmic C-terminus is essential for GPCR signaling. Previous work showed the LPHN C-terminus is necessary for synaptic function^31^ and that it interacts with postsynaptic SHANKs. Our data suggest that the LPHN intracellular tail likely functions as a multipurpose scaffold that regulates G protein and arrestin-3 interactions while also nucleating postsynaptic scaffolds^56^. We hypothesize that C-terminal residues can be extensively phosphorylated and this may lead to a bias toward arrestin-3. Deletion of the entire C-terminal tail completely abolished arrestin-3 binding. However, in contrast to the LPHN2 ΔCt mutant, the LPHN2 splice variants we used, with the exception of LPHN2v13, demonstrated TA-dependent arrestin-3 binding, albeit to differing levels. Together with the LPHN2 ΔCt results, this suggests that the membrane proximal tail sequence shared by all variants is essential for the TA-dependent recruitment of arrestin-3. Notably, the degree of that recruitment differed among variants, indicating that the precise sequence of the C-terminus determines the level of binding. The ability to recruit arrestin-3 did not correlate with Gα12/13 activation, suggesting that arrestin-3 binding to the LPHN2 does not depend on the Gα12/13 activation or the presence of Gα proteins. Thus, it is likely to serve both desensitization and signaling functions.

Recent work demonstrated that Gα12/13 engages ICL2 and ICL3 of LPHN2^43^. We confirmed the elimination of TA-dependent Gα12/13 activation in the LPHN2 AA mutant (ICL2)^43^, and showed that the ICL3 insertion in LPHN2v13 reduces Gα12/13 activation. We further showed that segments of the C-terminal tail also modulate G protein interactions, with the strongest effect observed in LPHN2v4 and v7. LPHN2v4 displays almost non-existent TA-dependent Gα13 activation, yet the most robust arrestin-3 recruitment. Conversely, LPHN2v3 activates Gα12/13 like LPHN2v1, yet exhibits virtually no arrestin-3 recruitment. This lack of correlation suggests that the LPHN2 C-terminal tail sequence shifts the bias of the receptor. Future studies will be required to interrogate the cell type specific distribution of different splice variants, level of their expression, and contribution to cell type specific LPHN2 signaling *in vivo*. Different variants may be selectively expressed in certain cell types in the brain, where their bias would lead to different outcomes.

The phosphorylation of GPCR cytoplasmic residues by GRKs precedes arrestin binding, and the phosphorylation patterns are believed to serve as code for the events that follow^40,49^. We found that GRK2/3/5/6 play a major role in arrestin-3 recruitment to LPHN2, but loss of these kinases does not completely abolish recruitment. Thus, other kinases are likely involved, and it would be of interest to explore their role, particularly of kinases with critical synaptic relevance: PKA, PKC and CamKII. Moreover, certain LPHN2 variants demonstrated elevated arrestin-3 recruitment in the presence of GRK2, while others were GRK2-insensitive. Surprisingly, the GRK2-dependent variants also exhibited accelerated kinetics of arrestin-3 binding, suggesting that these variants might have distinct functionality. Interestingly, variants sharing the relatively long C-terminal tail harboring a PDZ binding motif are all GRK2-independent, suggesting that this long tail may preclude GRK2 activation and/or contain sites for different kinases. Examining the interplay between aGPCR activation mechanism, kinase recruitment, phosphorylation barcodes, and the effects of all these factors on G protein vs arrestin-3 bias will advance our knowledge of aGPCR signaling and function.

## Methods

### KEY RESOURCE TABLE

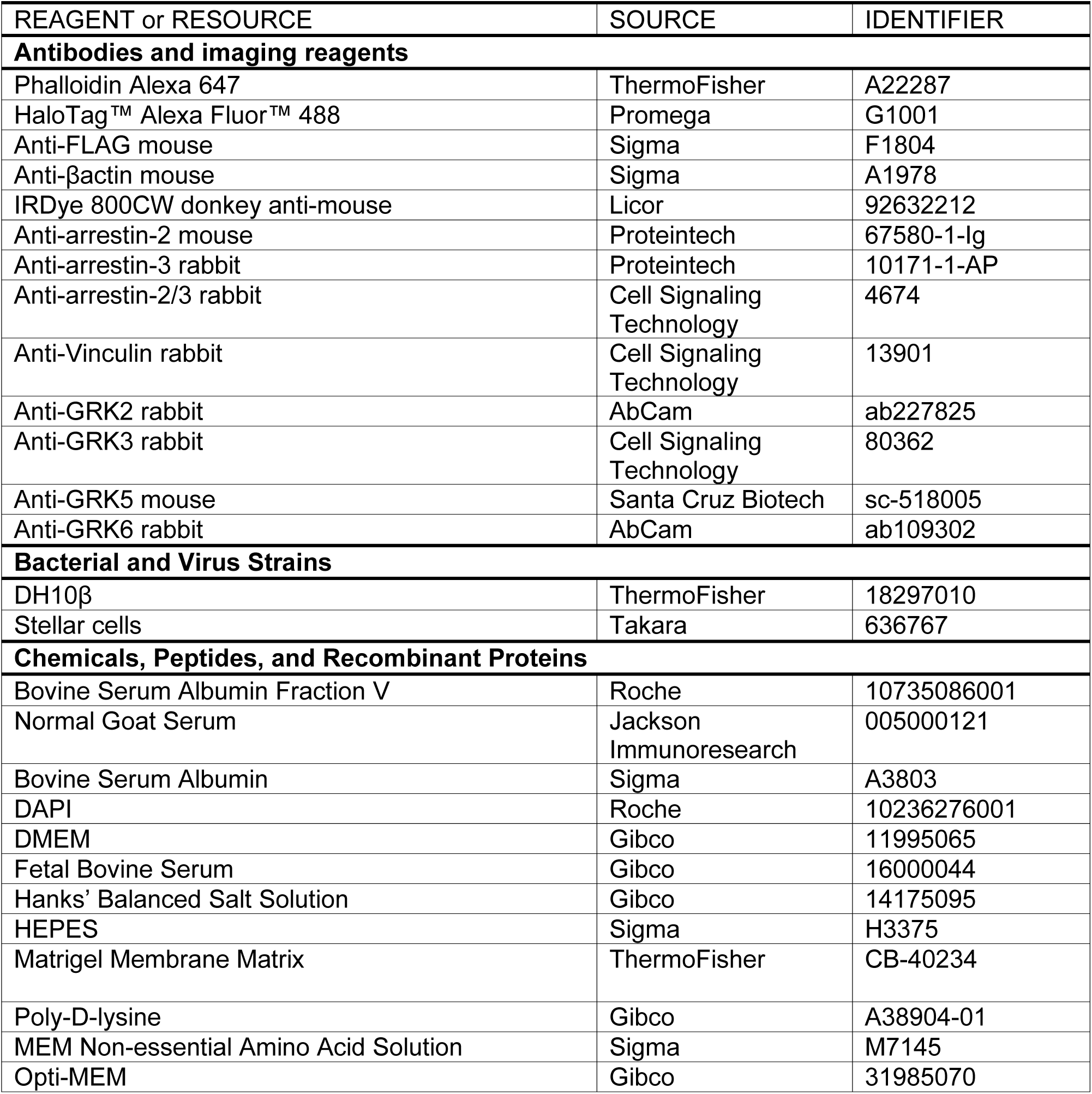
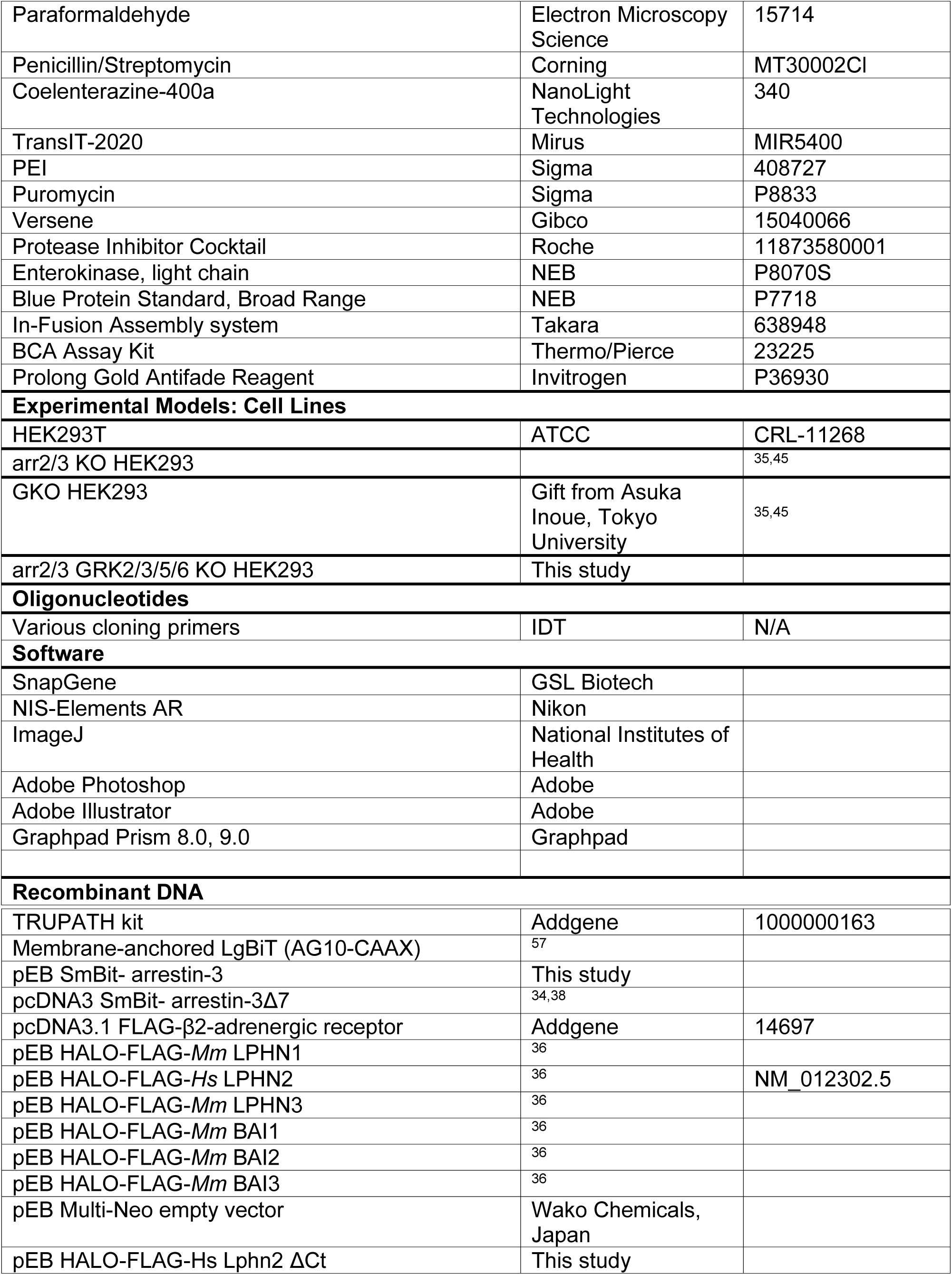
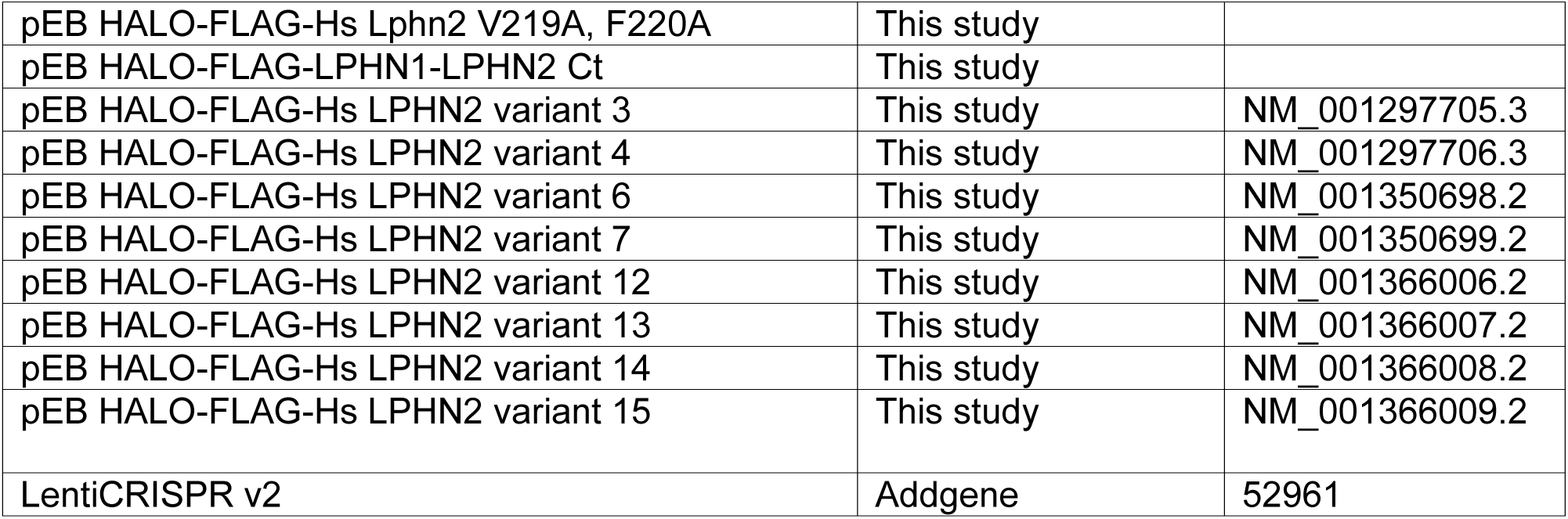

| REAGENT or RESOURCE | SOURCE | IDENTIFIER |
| --- | --- | --- |
| **Antibodies and imaging reagents** |  |  |
| Phalloidin Alexa 647 | ThermoFisher | A22287 |
| HaloTag™ Alexa Fluor™ 488 | Promega | G1001 |
| Anti-FLAG mouse | Sigma | F1804 |
| Anti-βactin mouse | Sigma | A1978 |
| IRDye 800CW donkey anti-mouse | Licor | 92632212 |
| Anti-arrestin-2 mouse | Proteintech | 67580-1-Ig |
| Anti-arrestin-3 rabbit | Proteintech | 10171-1-AP |
| Anti-arrestin-2/3 rabbit | Cell Signaling Technology | 4674 |
| Anti-Vinculin rabbit | Cell Signaling Technology | 13901 |
| Anti-GRK2 rabbit | AbCam | ab227825 |
| Anti-GRK3 rabbit | Cell Signaling Technology | 80362 |
| Anti-GRK5 mouse | Santa Cruz Biotech | sc-518005 |
| Anti-GRK6 rabbit | AbCam | ab109302 |
| **Bacterial and Virus Strains** |  |  |
| DH10β | ThermoFisher | 18297010 |
| Stellar cells | Takara | 636767 |
| **Chemicals, Peptides, and Recombinant Proteins** |  |  |
| Bovine Serum Albumin Fraction V | Roche | 10735086001 |
| Normal Goat Serum | Jackson Immunoresearch | 005000121 |
| Bovine Serum Albumin | Sigma | A3803 |
| DAPI | Roche | 10236276001 |
| DMEM | Gibco | 11995065 |
| Fetal Bovine Serum | Gibco | 16000044 |
| Hanks' Balanced Salt Solution | Gibco | 14175095 |
| HEPES | Sigma | H3375 |
| Matrigel Membrane Matrix | ThermoFisher | CB-40234 |
| Poly-D-lysine | Gibco | A38904-01 |
| MEM Non-essential Amino Acid Solution | Sigma | M7145 |
| Opti-MEM | Gibco | 31985070 |
| Paraformaldehyde | Electron Microscopy Science | 15714 |
| Penicillin/Streptomycin | Corning | MT30002CI |
| Coelenterazine-400a | NanoLight Technologies | 340 |
| TransIT-2020 | Mirus | MIR5400 |
| PEI | Sigma | 408727 |
| Puromycin | Sigma | P8833 |
| Versene | Gibco | 15040066 |
| Protease Inhibitor Cocktail | Roche | 11873580001 |
| Enterokinase, light chain | NEB | P8070S |
| Blue Protein Standard, Broad Range | NEB | P7718 |
| In-Fusion Assembly system | Takara | 638948 |
| BCA Assay Kit | Thermo/Pierce | 23225 |
| Prolong Gold Antifade Reagent | Invitrogen | P36930 |
| **Experimental Models: Cell Lines** |  |  |
| HEK293T | ATCC | CRL-11268 |
| arr2/3 KO HEK293 |  | 35,45 |
| GKO HEK293 | Gift from Asuka Inoue, Tokyo University | 35,45 |
| arr2/3 GRK2/3/5/6 KO HEK293 | This study |  |
| **Oligonucleotides** |  |  |
| Various cloning primers | IDT | N/A |
| **Software** |  |  |
| SnapGene | GSL Biotech |  |
| NIS-Elements AR | Nikon |  |
| ImageJ | National Institutes of Health |  |
| Adobe Photoshop | Adobe |  |
| Adobe Illustrator | Adobe |  |
| Graphpad Prism 8.0, 9.0 | Graphpad |  |
| **Recombinant DNA** |  |  |
| TRUPATH kit | Addgene | 1000000163 |
| Membrane-anchored LgBiT (AG10-CAAX) | <sup>57</sup> |  |
| pEB SmBit- arrestin-3 | This study |  |
| pcDNA3 SmBit- arrestin-3Δ7 | <sup>34,38</sup> |  |
| pcDNA3.1 FLAG-β2-adrenergic receptor | Addgene | 14697 |
| pEB HALO-FLAG-*Mm* LPHN1 | <sup>36</sup> |  |
| pEB HALO-FLAG-*Hs* LPHN2 | <sup>36</sup> | NM_012302.5 |
| pEB HALO-FLAG-*Mm* LPHN3 | <sup>36</sup> |  |
| pEB HALO-FLAG-*Mm* BAI1 | <sup>36</sup> |  |
| pEB HALO-FLAG-*Mm* BAI2 | <sup>36</sup> |  |
| pEB HALO-FLAG-*Mm* BAI3 | <sup>36</sup> |  |
| pEB Multi-Neo empty vector | Wako Chemicals, Japan |  |
| pEB HALO-FLAG-*Hs* Lphn2 ΔCt | This study |  |
| pEB HALO-FLAG-Hs Lphn2 V219A, F220A | This study |  |
| pEB HALO-FLAG-LPHN1-LPHN2 Ct | This study |  |
| pEB HALO-FLAG-Hs LPHN2 variant 3 | This study | NM_001297705.3 |
| pEB HALO-FLAG-Hs LPHN2 variant 4 | This study | NM_001297706.3 |
| pEB HALO-FLAG-Hs LPHN2 variant 6 | This study | NM_001350698.2 |
| pEB HALO-FLAG-Hs LPHN2 variant 7 | This study | NM_001350699.2 |
| pEB HALO-FLAG-Hs LPHN2 variant 12 | This study | NM_001366006.2 |
| pEB HALO-FLAG-Hs LPHN2 variant 13 | This study | NM_001366007.2 |
| pEB HALO-FLAG-Hs LPHN2 variant 14 | This study | NM_001366008.2 |
| pEB HALO-FLAG-Hs LPHN2 variant 15 | This study | NM_001366009.2 |
| LentiCRISPR v2 | Addgene | 52961 |

## RESOURCE AVAILABILITY

### Lead contact

Further information and requests for resources and reagents should be directed to and will be fulfilled by the lead contact, Richard C. Sando.

### Materials availability

All materials generated in this study will be openly shared upon request.

## EXPERIMENTAL MODELS AND SUBJECT DETAILS

### Cell Lines

HEK293T cells (ATCC # CRL-11268) were maintained in DMEM (Gibco Cat# 11995065) containing 10% FBS (Gibco Cat# 16000044), 1X Penicillin-Streptomycin (Corning Cat# MT30002Cl) at 37°C and 5% CO_2_ for a maximum of 25 passage numbers. HEK293T cells were used to examine protein expression and cell surface localization. HEK293 arr2/3 KO used for arrestin-3 assays were originally generated by Milligan lab from University of Glasgow ^35^ and were a gift from Drs. Vsevolod Gurevich and Chen Zheng (Vanderbilt University). The HEK293 arr2/3 GRK2/3/5/6 KO line was used for discerning the role of GRKs on arrestin-3 binding to GPCRs. See section below describing the generation of this line. GKO HEK293 cells used for all BRET2 assays were originally a gift from Asuka Inoue (Tokyo University, Japan) ^35,45^ and were provided to our studies as a gift from Drs. Vsevolod Gurevich and Chen Zheng. GKO HEK293 cells were maintained in DMEM (Gibco Cat# 11995065) containing 10% FBS (Gibco Cat# 16000044), 1X Penicillin-Streptomycin (Corning Cat# MT30002Cl), and 1x MEM Non-essential Amino Acid (NeA) Solution (Sigma Cat# M7145) at 37°C and 5% CO_2_.

### CRISPR/Cas9-mediated knockout of arrestin-2/3 in HEK293 cells

Stable HEK293 arrestin-2/3 knockout cells were generated by transient transfection using self-made PEI reagent (Sigma-Aldrich, 408727, diluted to 10 μg/ml, pH 7.2, adjusted with HCl) of the CRISPR Control or ΔQ-GRK (GRK2/3/5/6-KO) cells^47^ with lentiCRISPR v2 plasmid (Addgene #52961) containing target-specific gRNAs (**Table 1**). To produce these vectors, the complementary forward and reverse oligo nucleotides listed in Table 1 were annealed and ligated into BsmBI-restricted lenti-CRISPR v2 vector. Specific knockout of arrestin-2 or 3 isoforms was achieved by simultaneous transfection of the respective four gRNA constructs, double knockout of arrestin-2/3, by transfection of all eight gRNA constructs. Selection of transfected cells was performed using 1 µg/ml puromycin (Sigma-Aldrich #P8833) and subsequently single-cell clones were established. Absence of arrestin-2 and 3 was confirmed by western blot using isoform specific antibodies (arrestin-2 Proteintech 67580-1-Ig, arrestin-3 Proteintech 10171-1-AP, and arrestin-2/3 Cell Signaling Technology #4674).

**Table 1:**
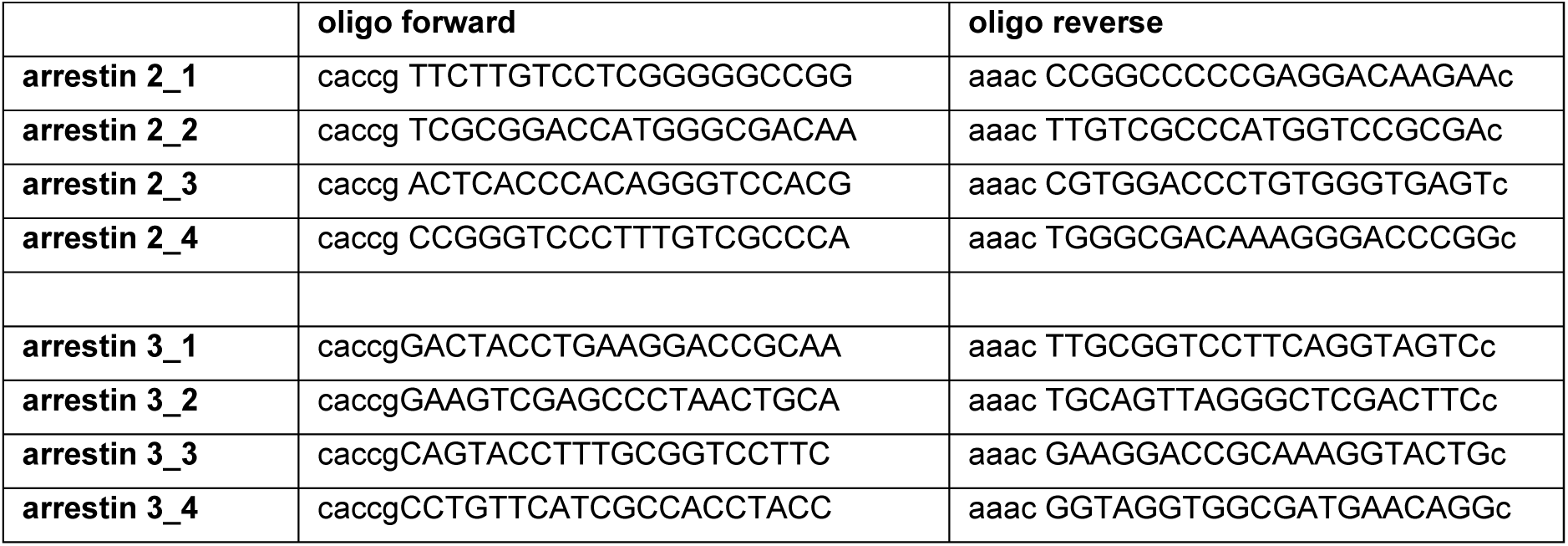
gRNA sequences for targeting arrestin-2 or 3. Complementary forward and reverse oligos were annealed and ligated into the BsmBI-restricted lentiCRISPR v2 vector^58^ (Addgene #52961). gRNA sequence is shown in capital letters, overhangs for ligation in lowercase letters. Sequences of gRNAs were validated in^58^.

### Plasmids

All aGPCR overexpression cDNAs used were encoded in the pEB Multi-Neo vector (Wako Chemicals, Japan). The HALO-FLAG-aGPCR constructs contained the following sequence of features fused upstream of the native tethered agonist sequence: IgK signal peptide (METDTLLLWVLLLWVPGSTGDAGAQ) – HALO-tag – linker (GGSGGSGGS) – FLAG (DYKDDDDK). Enterokinase-mediated cleavage following DDDDK/ was designed to expose the native TA region for the aGPCR: LPHN1 (N-TNFAVLMAH…-C), LPHN2 (N-TNFAILMAH…-C), LPHN3 (N-TNFAVLMAH…-C), BAI1 (N-STFAILA…-C), BAI2 (N-STFAVLAQ…-C), and BAI3 (N-STFAILAQ…-C). See Resource Table for information on human Lphn2 transcript variants. The TRUPATH kit was purchased from Addgene (#1000000163). Cells transfected with empty pEB-Multi-Neo were used as negative control conditions in all experiments. All molecular cloning was conducted with the In-Fusion Assembly system (Takara #638948) and NEB HiFi DNA Assembly master mix (NEB E2621). For arrestin-3 membrane recruitment NanoBiT assays, the indirect approach using membrane-tethered LgBit (AG10-CAXX LargeBit) and SmBit-arrestin-3 was used to minimize the impact of fusions to the C-terminus of aGPCRs.

### Antibodies

The following antibodies and reagents were used at the indicated concentrations: anti-β-actin mouse (Sigma, A1978, 1:10,000); anti-FLAG mouse (Sigma, F1804, 1:2,000); Alexa Fluor 647 Phalloidin (Invitrogen Cat# A22287; 1:40 diluted in methanol), HaloTag™ Alexa Fluor™ 488 (Promega, G1001, 1 µM final concentration), IRDye 800CW donkey anti-mouse (Licor, 92632212, 1:10,000), anti-β-arrestin 1 mouse (Proteintech 67580-1-Ig, 1:1,000); anti-β-arrestin 2 rabbit (Proteintech 10171-1-AP, 1:1,000); anti-β-arrestin 1/2 rabbit (Cell Signaling Technology #4674, 1:1,000); anti-Vinculin (Cell Signaling Technologies #13901, 1:1,000); anti-GRK2 rabbit (AbCam #ab227825, 1:1,000); anti-GRK3 rabbit (Cell Signaling Technologies #80362, 1:250); anti-GRK5 mouse (Santa Cruz Biotechnologies #sc-518005, 1:250); and anti-GRK6 rabbit (AbCam #ab109302, 1:1,000).

### NanoBiT Arrestin Assay

The arrestin-3 assay was performed using the NanoBiT membrane recruitment system ^57^. HEK293 arr2/3 KO^35^ and arr2/3 GRK-2/3/5/6 KO cells were plated into 12-well plate at a density of 4-5 × 10^5^ cells and 1×10^6^ respectively in 1 mL per well in 1x DMEM (Gibco Cat# 11995065) plus 10% FBS (Gibco Cat# 16000044) and 1X Penicillin-Streptomycin (Corning Cat# MT30002Cl). After 20-24 hours, cells were co-transfected via TransIT-2020 (Mirus MIR5400) with receptor of interest (0.3 µg/µL); membrane-anchored AG10-CAXX LargeBit (0.05 µg/µL) and SmBiT-arrestin3 (0.05 µg/µL) for a total of 0.4 µg DNA per transfection condition. When GRK2 contribution was tested 1 µL of GRK2 (0.3 µg/µL) was added and the negative control was balanced with the addition of pEB (0.3 µg/µL). Each condition required 96 µL of room-temperature 1x Opti-MEM (Gibco Cat# 31985070), 1 µL each DNA plasmid at specified concentrations, and 3 µL of room-temperature and gently-vortexed TransIT-2020 reagent. The TransIT-2020:DNA complexes were mixed via pipetting 10 times and incubated at room temperature for 20 mins before adding drop-wise in the well. The plate was rocked gently from side to side and incubated at 37°C 24 hours before harvesting. From each well, media was aspirated, and cells were washed with 1 mL warm PBS. Cells were then detached with 300 µL warm Versene (Gibco Cat# 15040066) for 5 min at 37°C, and then transferred to 1.5 mL Eppendorf tubes with 1 mL DMEM. Cells were then plated at 200 µL per well in Matrigel-coated 96-well white assay plate. Twenty-four hours after plating, media was aspirated and replaced with 80 µL of HBSS + 20 mM HEPES+ 5 mM glucose. Ten µL of 100 µM Coelenterazine-400a (NanoLight Technologies Cat# 340) in PBS was added to each well before the measurements of total luminescence were initiated in a repeated manner. After 10 cycles of measurement (approximately 10 min), 10 µl of receptor activators were added (Enterokinase (NEB, P8070L) to 0.055 U final concentration for Lphn1-3 and Bai1-3, and isoproterenol (Sigma Cat# I6504) to 1 µM final concentration for β2AR, or PBS as a negative control) and the repeated measurements were resumed for an additional 60 minutes. A ratio was calculated from the luminescence with activator and without (PBS) and then normalized to the average of 5 basal reads before addition of activator. For the Lphn2 variants, normalization was also performed based on the surface expression level of each variant.

### TRUPATH BRET2 assays

HEK293 G protein K.O. (GKO) cells were plated into 12-well plates at a density of 4-4.5 × 10^5^ cells in 1 mL per well. HEK293 GKO media contained 1x DMEM (Gibco Cat# 11995065) plus 10% FBS (Gibco Cat#16000044), 1X Penicillin-Streptomycin (Corning Cat#MT30002Cl) with 1x MEM Non-essential Amino Acid (NeA) Solution (Sigma Cat# M7145). After 20-24 hours, cells were co-transfected with the receptor-of-interest and TRUPATH plasmids at 1:1:1:1 DNA ratio (receptor:Gα-RLuc8:Gβ:Gγ-GFP2) via TransIT-2020 (Mirus Cat# MIR5400). Each condition required 97 µL of room-temperature 1x Opti-MEM (Gibco Cat# 31985070), 1 µL each DNA plasmid at 1 µg/µL concentration, and 3 µL of room-temperature and gently-vortexed TransIT-2020 reagent. The TransIT-2020:DNA complexes mixture were gently mixed via pipetting 10 times and incubated at room temperature for 20 mins before adding drop-wise in the well. The plate was rocked gently side to side and incubated at 37°C 24 hrs before harvesting. In each well, media was aspirated after 24 hrs, and cells were washed with 1 mL warm PBS. Cells were detached with 300 µL warm Versene (Gibco Cat# 15040066) and incubated at 37°C for 5 mins, resuspended via pipetting 10 times, and then transferred to 1.5 mL Eppendorf tube with 1 mL DMEM containing 1x NeA. Cells were then plated at 200 µL per well in Matrigel-coated 96-well white assay plate. Each experimental condition was plated into six separate wells within the 96-well assay plate. BRET2 assays were performed 48 hrs after transfection. In each well, media was aspirated and cells were incubated in 80 µL of 1x Hanks’ balanced Salt Solution (Gibco Cat# 14175095) with 20 mM HEPES (Sigma Cat# H3375, pH 7.4) and 10 µL 100 µM Coelenterazine-400a (NanoLight Technologies Cat# 340) which was reconstituted in PBS. Receptors were activated by addition of 10 µL of enterokinase enzyme/agonist in PBS to achieve the following final concentrations: LPHN1-3 and BAI1-3 with 0.055 U of Enterokinase (NEB, P8070L); β2AR with 1 µM of isoproterenol (Sigma Cat# I6504). BRET intensities were measured via BERTHOLD TriStar2 LB 942 Multimode Reader with Deep Blue C filter (410nm) and GFP2 filter (515 nm). The BRET ratio was obtained by calculating the ratio of GFP2 signal to Deep Blue C signal per well. The BRET2 ratio of the three wells per condition were then averaged. Net BRET2 was subsequently calculated by subtracting the BRET2 ratio of cells expressing donor only (Gα-RLuc8) from the BRET2 ratio of each respective experimental condition. Net BRET2 differences were then compared as described in the Figures: Basal, by subtracting Net BRET2 ratios of empty vector (EV) from Receptor; TA-exposed, by subtracting Net BRET2 ratios of vehicle from activator-treated. When repeated measurements were carried out, the media was aspirated and replaced with 80 µl of HBSS +20 mM HEPES+ 5 mM glucose. Ten µL 100 µM Coelenterazine-400a (NanoLight Technologies Cat# 340) in PBS was added to each well before the measurements were initiated in a repeated manner. After 5-10 cycles of measurement (approximately 10 min), 10 µl of receptor activators were added (Enterokinase (NEB, P8070L) to 0.055 U final concentration, and the repeated measurements were resumed for additional 60 minutes. The ΔNet BRET2 calculated for each receptor (EK-PBS) was normalized to the average of 5 basal reads before addition of activator. For the Lphn2 variants, additional normalization was performed based on the surface expression level of each variant.

### Surface labeling and immunocytochemistry

Cover glass (#0, 12 mm, Carolina Biological Supply Company #633009) was placed into 24-well plates and coated for 2 hrs with 100 µL of 50 µg/mL poly-D-lysine (Gibco #A38904-01) in the 37°C tissue culture incubator. Excess poly-D-lysine was removed, coverslips were washed 3x with sterile ddH_2_O and dried for 30 mins. HEK293T cells were plated at 1.5-2 × 10^5^ cells/well in 0.5 mL complete DMEM. After 16-24 hrs, cells were transfected with indicated experimental plasmid via TransIT-2020 (Mirus MIR5400) with a total of 0.5 µg DNA amount/condition/well. After 48 hrs post-transfection, HALO-Tag AlexaFluor488 (Promega, #G1001) was applied at 1 µM for 1 hr at 37°C, prior to being fixed. Cells were then washed briefly once with PBS, fixed with 4% PFA (Electron Microscopy Science Cat# 15714)/4% sucrose/PBS for 20 min at 4°C, and washed 3 x 5 mins in PBS. Fluorescently conjugated Alexa Fluor 647 Phalloidin (Invitrogen Cat# A22287; 1:40 diluted in methanol) and DAPI (Sigma Cat# 10236276001; 1:1,000) diluted in PBS was applied for 30 mins. Cells were then washed 2x times in PBS, and mounted on UltraClear microscope slides (Denville Scientific Cat# M1021) using 10 µL ProLong Gold antifade reagent (Invitrogen, #P36930) per coverslip. Low-magnification images were collected with a 20x objective and high-magnification images a 60x objective (see Imaging section for details). Surface expression levels were estimated by the ratio of sum intensity of channel 488 for surface HALO versus the sum intensity of channel 647 for Phalloidin from the 20x images after rolling ball background subtraction set as 19.75 µm.

### Confocal imaging of fixed samples

Images were acquired using a Nikon A1r resonant scanning Eclipse Ti2 HD25 confocal microscope with a 20x (Nikon #MRD00205, CFI60 Plan Apochromat Lambda, N.A. 0.75), and 60x (Nikon #MRD01605, CFI60 Plan Apochromat Lambda, N.A. 1.4) objectives, operated by NIS-Elements AR v4.5 acquisition software. Laser intensities and acquisition settings were established for individual channels and applied to entire experiments. Image analysis was conducted using Nikon Elements and ImageJ. Brightness was adjusted uniformly across all pixels for a given experiment for figure visualization purposes. Quantification of fluorescence intensities was conducted by imaging 3-5 image frames per biological replicate, which were averaged to generate a single biological replicate value. The averaged value for each replicate is depicted as open circles in each graph.

### Immunoblotting

HEK293T were plated in 12-well plates at 450,000 cells/well density. Cells were transfected 24 hrs later via TransIT-2020 (Mirus MIR5400) at 1 µg DNA total/well. After 48-hrs post-transfection, cells were briefly washed 1X with PBS, and samples were collected in 0.75 mL/well 1X RIPA buffer (final concentration, 50 mM Tris pH7.6, 150 mM NaCl, 1% NP-40, 0.5% sodium deoxycholate, 0.1% SDS) containing protease inhibitors (Roche Cat# 11873580001) for 1 hr at 4°C with gentle rotation. Homogenate was centrifuged for 15 mins at 14,000xg at 4°C, the supernatant was collected and total protein was measured via BCA assay (Thermo, Pierce, #23225). Samples were then diluted in sample buffer containing 312.5 mM Tris-HCl pH 6.8, 10% SDS, 50% glycerol, 12.5% 2-mercaptoethanol, bromophenol blue, and protease inhibitors (Roche Cat# 11873580001) and 3 µg total protein/well run on 4-20% SDS-PAGE gels (BioRad mini-protean TGX Cat#4561096) at 75 V at room temperature. The Blue Protein Standard, Broad Range (NEB Cat# P7718) was used as a protein molecular weight ladder. Protein was transferred onto nitrocellulose transfer membrane in transfer buffer (25.1 mM Tris, 192 mM glycine, 20% methanol) at 100 V for 2 hrs at 4°C. Membranes were blocked in 4% bovine serum albumin (BSA, Sigma Cat# 10735086001) /TBST (20 mM Tris-HCl pH 7.4, 150 mM NaCl, 0.05% Tween-20) for 1 hr at room temperature, incubated in primary antibodies diluted into 4% BSA/TBST overnight at 4°C (anti-FLAG mouse 1:2,000; anti-βactin mouse 1:10,000), washed 3 x 5 mins in TBST, incubated in corresponding secondary antibodies (Licor IRDye 800CW donkey anti-mouse Cat#92632212) diluted 1:10,000 into TBST, washed 5 x 5 mins in TBST, and imaged on a Licor Odyssey system.

## QUANTIFICATION AND STATISTICAL ANALYSIS

### Statistics

All data are expressed as means ± SEM and represent the results of at least three independent biological replicates, as indicated within each Figure Legend and as open circles within bar graphs. Statistical significance was determined using the two-tailed Student’s t-test, one-way ANOVA with following *post hoc* Tukey tests for multiple comparisons, or two-way ANOVA with following *post hoc* Tukey tests for multiple comparisons, as indicated in the Figure Legends. Data analysis and statistics were performed with Microsoft Excel, GraphPad Prism 8.0 and GraphPad Prism 9.0.

## Acknowledgements

We thank all members of the Sando laboratory for critical feedback on the study. This study was supported by grants from the NIH (R00-MH117235 and DP2MH1401324 to RS) and Alfred Sloan Foundation (Sloan Fellowship in Neuroscience to RS).

## Author Contributions

K. Garbett performed all BRET2 and NanoLuc experiments. K. Garbett and R. Sando performed all molecular cloning. R. Sando conducted immunoblotting and imaging experiments. K. Garbett and R. Sando performed data analysis and interpretation. K. Garbett and R. Sando designed the study with the critical help of C. Zheng and V.V. Gurevich. J. Drube and C. Hoffmann generated and characterized the arr2/3 GRK2/3/5/6 KO and CRISPR control cell lines. K. Garbett and R. Sando wrote the manuscript, and all authors edited it.

