## Supplemental Figures for "Cytoplasmic tail composition modulates the G protein and arrestin-3 signaling bias of the adhesion GPCR LPHN2"

**SUPPLEMENTAL DATA FIGURES and FIGURE LEGENDS**

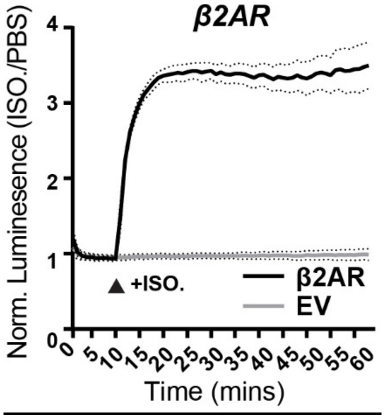


Figure S1: **arrestin-3 recruitment to β2-adrenergic receptor (β2AR)**

Time course of arrestin-3 recruitment in cells transfected with β2AR or empty vector (EV) treated with 1 μM Isoproterenol (ISO) or PBS.

Numerical data are means ± SEM from 3 independent biological replicates.

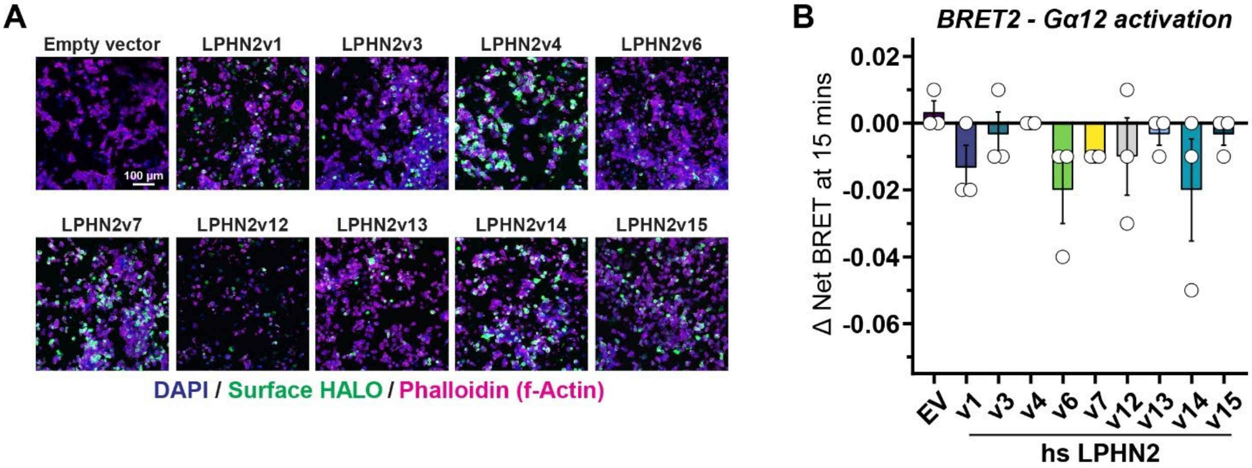


Figure S2: **Additional characterization of LPHN2 transcript variants**

**A,** low-magnification images of HEK293T cells transfected with the indicated conditions, live labeled for surface HALO488, and subsequently labeled for Phalloidin (f-Actin) and DAPI.

**B,** ΔNet BRET2 measurements using the Gα12 TRUPATH BRET2 sensor at 15-min post-EK treatment with indicated human Lphn2 variants compared to empty vector control.

Numerical data are means ± SEM from 3 independent biological replicates (depicted as open circles).

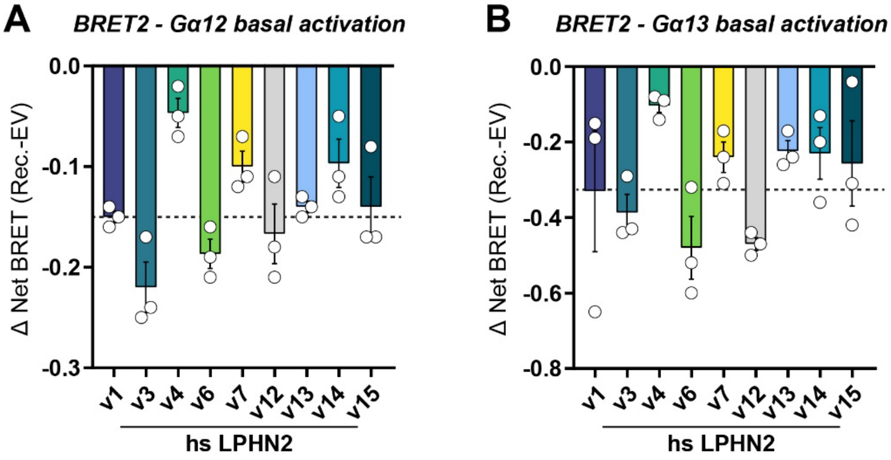


<u>Figure S3:</u> **Basal Gα12/13 activation by LPHN2 transcript variants**

**A,** TRUPATH Gα12 basal activation (before addition of EK) measured by Δ Net BRET (Receptor vs empty vector (EV)).

**B,** similar to **A**, using Gα13 TRUPATH sensor.

Numerical data are means ± SEM from 3 independent biological replicates (depicted as open circles).

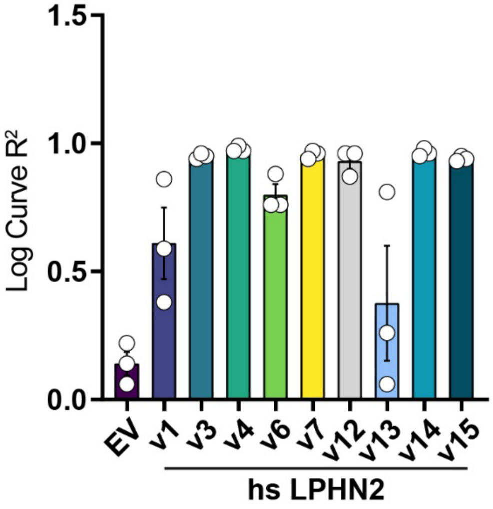


<u>Figure S4:</u> **Logarithmic fit of the curves of arrestin-3 recruitment obtained with LPHN2 variants shown in Figure 6**

$R^2$ values from fitting indicated experimental conditions to a logarithmic curve.

Numerical data are means ± SEM from 3 independent biological replicates (depicted as open circles).

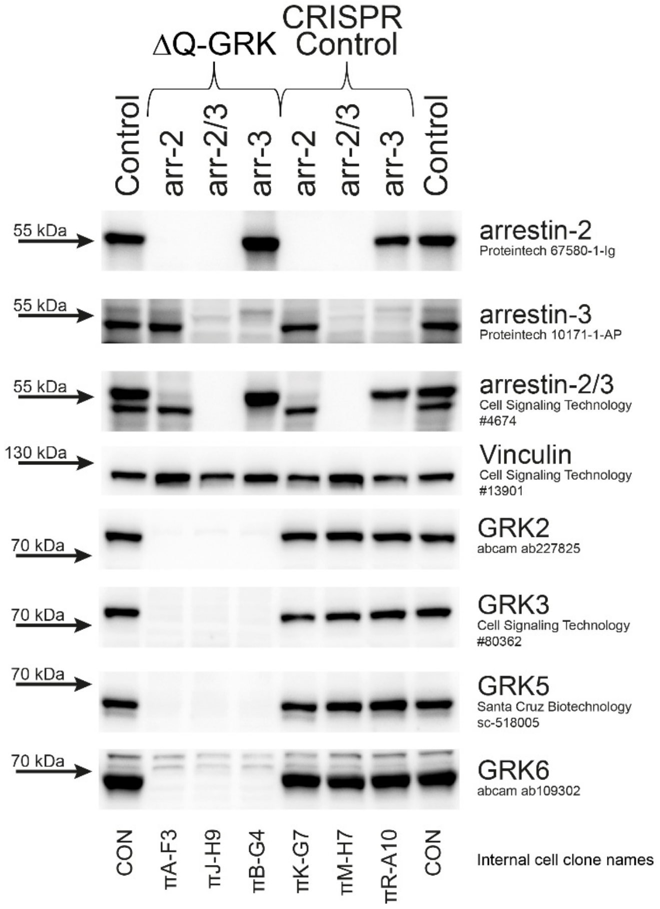


Figure S5: **Generation and validation of arr-2/3 CRISPR control and arr-2/3 GRK2/3/5/6 KO HEK293 cell lines**

Immunoblots validating CRISPR/Cas9-mediated deletion of arr-2/3 from ΔQ-GRK (GRK2/3/5/6) and CRISPR control lines. Vinculin was used as a loading control. See Methods section for details regarding the generation of these cell lines.
